# BREAKTHROUGH REPORT - Large scale measurement of protein synthesis rates over the diurnal cycle reveal pathway specific regulation of translation

**DOI:** 10.1101/2021.03.15.435498

**Authors:** Owen Duncan, A. Harvey Millar

## Abstract

Plants have a diurnal separation of metabolic fluxes and a need for differential maintenance of protein machinery in the day and night. To directly assess aggregate protein translation and degradation for specific proteins and to estimate the ATP investment involved, the individual rates of synthesis and degradation of hundreds of different proteins were measured in *Arabidopsis thaliana* rosettes. This quantification of translation control through incorporation of heavy hydrogen into newly synthesised protein confirmed that most protein synthesis occurs during the day hours (~3:1 day:night), but revealed it was highly divergent across functional categories. Proteins involved in photosynthesis, especially components of the light harvesting complexes, were synthesized much faster in the day (~10:1), while the protein components of carbon metabolism and vesicle trafficking were translated at similar rates day or night. Comparison of aggregate translation with a range of comparable studies using polysomal loading of transcripts or transcript abundance suggests that translational control is a major contributor to effective gene expression over the diurnal cycle. Diurnal protein degradation rate was observed to be tightly coordinated with protein synthesis rate as few leaf proteins changed in abundance despite reduced translation rates during the night. The direct, quantitative and aggregate analysis of protein synthesis provides an integration of transcriptional, post-transcriptional and translational control of leaf proteome homeostasis and an opportunity to assess the ATP investments involved. The data reveals how the pausing of photosystem synthesis and degradation at night allows the redirection of a decreased energy budget to a selective night-time maintenance schedule.

**One sentence summary:** Advances in collection and processing of stable isotope incorporation data has allowed characterisation of differential protein synthesis rates over the diurnal cycle which reveal both global and specific translational regulation programs of core biochemical pathways.

## Introduction

When, where and at what rate genetic information is expressed are important determinants of an organism’s success. Detailed information on some elements of this process in model organisms have been reported; e.g. genome sequences, genome modification status, transcriptional rates and abundances, protein abundances and post translational modifications. However, comparatively little is known about the key energetic commitment step in this process: the rate of translation of specific proteins. In comparison to transcription, translation is an expensive process; consuming at least an order of magnitude more ATP than transcription (Wagner 2005). In some developmental stages in plants, translation of all proteins accounts for up to 38% of the respiratory ATP available (Li et al. 2017). In mammalian systems which have comparatively stable energetic conditions, this cost is minimised predominantly through transcript driven control (Li, Bickel, and Biggin 2014). In plants however, there is substantial evidence that this is not the case; possibly due to strongly varying energy availability driven by photosynthesis in the day - night cycle. Instead, evidence points to much more protein synthesis occurring during the day hours than occurs during the night hours, despite evident, but numerically smaller fluctuations in transcript abundance or polysomal loading over the same cycle (Ishihara et al. 2015; Piques et al. 2009).

Recent studies in Arabidopsis have focused on the association of transcripts with ribosomes as a proxy measurement of translation of the encoded protein (Juntawong et al. 2014; Missra et al. 2015; Yamasaki et al. 2015). This is a powerful technique due to its general applicability, compatibility with mainstream nucleotide sequencing technologies and the depth of analysis yielded. However, as a proxy rather than a direct measurement; multiple levels of regulation can confound the correlation of these measurements with protein translation and therefore cellular function. For example, Missra et al. (2015) showed that transcripts encoding components of the cytosolic ribosome are in high translation states at night, while end of day and end of night ribosome abundance measurements show that they do not accumulate over this period (Pal et al. 2013). Given the degradation of ribosomal protein subunits is mostly uniform and slow (Salih et al. 2020; Li et al. 2017), this suggests that either fruitless and expensive synthesis/degradation cycles or regulation not captured by measurements of mRNA translation state was occurring (Missra et al. 2015).

Earlier, pre-’omics studies where both protein translation and polysomal association of transcripts were measured show that there is substantial diversity of regulation occurring both in the association of transcripts with ribosomes and with translation once initiated. Investigations into the regulation of CATALASE ISOZYME 2 (CAT2) in *Zea mays* demonstrated that while transcript abundance and polysomal association was equal in both the dark and the light, protein was only produced in the light (Skadsen and Scandalios 1987). Similarly, light responsive translational control of RUBISCO expression was demonstrated in *Amaranthus hypochondriacus* where translation preceded transcriptional regulation in dark to light transitions (Berry et al. 1986). Light based regulation of the translation of plastid FERREDOXIN-1 (Fed-1) is relatively well understood with the transcript only being associated with ribosomes in the light in the absence of photosynthetic flux inhibitors (Petracek et al. 1997) due to the presence of a 5’UTR located mRNA light response element which is capable of conferring light regulation on any associated open reading frame (Hansen et al. 2001). The understanding of these complex mechanisms of post transcriptional regulation have been built through measuring both the input and the output of the ribosome, not just one side of the equation.

The ability to investigate aggregate protein translation however has been hampered by the laborious nature of making translational measurements; most studies have involved incubation of plant tissues with radio-labelled Met followed by immuno-precipitation of the target protein and gel based separations followed by radio densitometry. These techniques are difficult for individual proteins and limited in scale, but have yielded insight into intricate mechanisms of post transcriptional regulation in some cases (Qi et al. 2011). Stable isotopes supplied in the form of inorganic compounds have been used to good effect to measure bulk protein synthesis rates in diurnal cycles (Ishihara et al. 2015) and individual protein synthesis rates over the course of days (Li et al. 2017; Nelson et al. 2014). Generally speaking, ^15^N labelling of plants works very inefficiently at night due to the predominance of ferredoxin linked nitrate reduction in the plastid as a means of N assimilation (Li et al. 2017; Nelson et al. 2014) and ^13^CO_2_ labelling is effectively absent at night due to the lack of photosynthesis. These considerations limit the ability of either label to readily capture an in-depth catalogue of night time effects on protein synthesis.

With these limitations in mind, we have harnessed the ability of ^2^H in water to be rapidly assimilated into the amino acid pool during both the day and the night and mass spectrometry based proteomics to provide time resolved, protein specific assessments of translation and degradation over the day / night cycle. This approach complements protein translation and degradation at the total pool level and gene specific measurements of polysomal association of mRNA to provide a more complete assessment of the translation process from the protein side. The aggregate nature of the data also allows the metabolic, developmental and physiological responses to the availability of energy to be incorporated into the interpretation of protein translation data. Energetically expensive processes that are only needed in the day may be best conducted in the light when energy is plentiful, while a reduced set of essential functions may need to be maintained or enhanced without exhausting starch supplies in the night.

## Results

### Determining Protein Synthesis and Degradation Rates in the Day and Night

Three week old *Arabidopsis thaliana* Col-0 plants were grown hydroponically on a 12 hour light / 12 hour dark diurnal cycle. Forty five minutes before lights on, day labelled plants were transferred in to heavy labelled (5% ^2^H_2_O / 95% H_2_O) Hoagland’s media. At lights off, whole rosettes were harvested, quenched in liquid N_2_ and stored as an end of day sample. Forty-five minutes prior to this, a second set of plants were transferred into 5% ^2^H_2_O and grown through the 12 hour dark period before sampling immediately before lights on, representing an end of night sample. Matching samples grown in 100% H_2_O Hoagland’s were collected at end of day and end of night. Proteins from these samples were extracted, digested and analysed by standard data dependent mass spectrometry workflows (Figure 1a).

**Figure 1.**
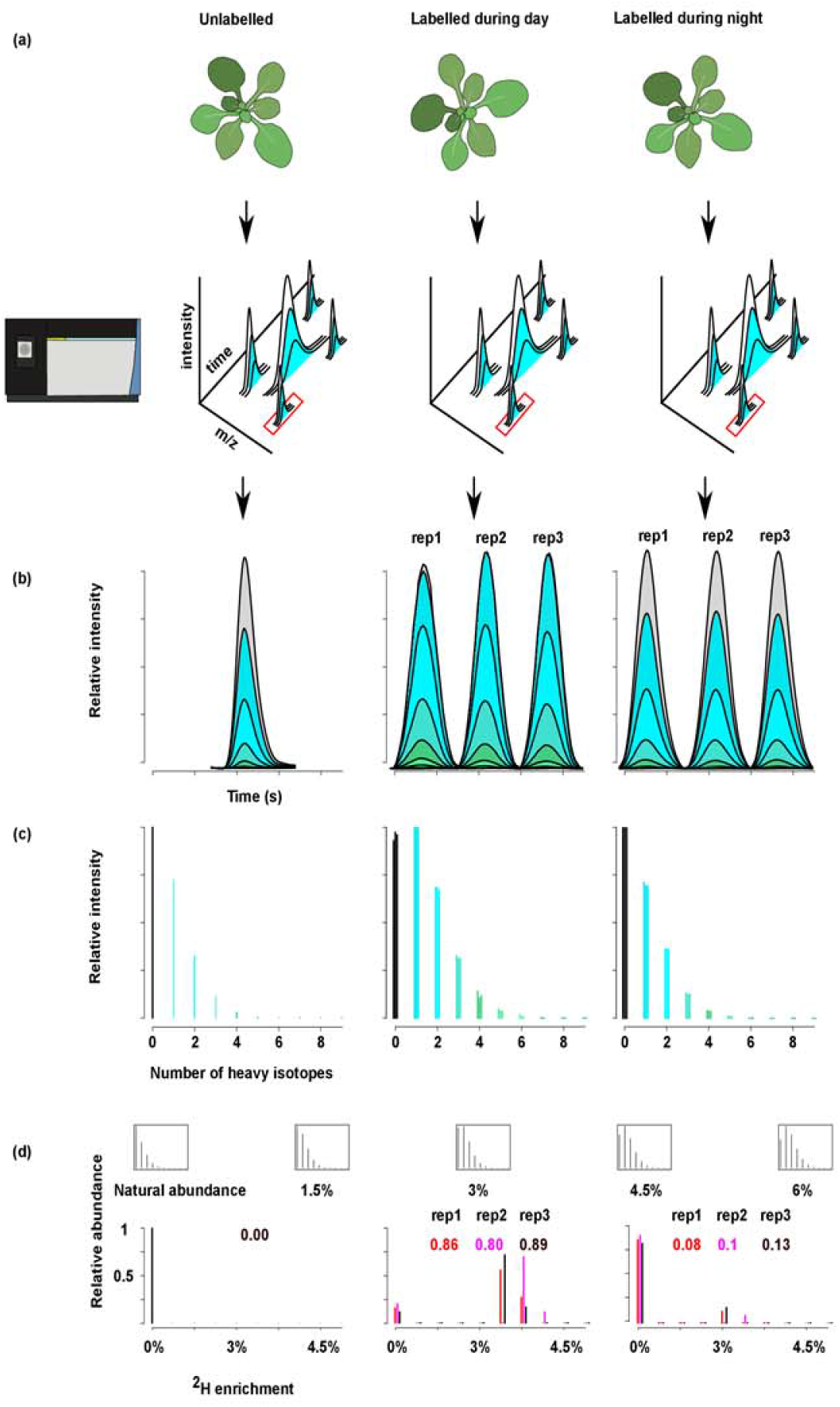
Labelling of Arabidopsis leaf proteins in the day and night for turnover rate calculation. (a) Approximately 216 *Arabidopsis thaliana* plants were grown in Hoagland’s media in l2h day / 12h night for 21 days. On the morning of the 21st day, three trays each containing 24 plants were transferred into media containing 5% ^2^H_2_O before sampling at the end of day. A second set of 72 plants were labelled through 12 hours of night while concurrently grown unlabelled plants were sampled at the end of day and end of night. Proteins were extracted, digested, fractionated and analysed by standard bottom-up data dependent mass spectrometry. (b) Labelled and unlabelled runs were retention time aligned and MS^1^ features with matching peptide identifications were cross-extracted between three replicates of day labelled plants and three replicates of night labelled plants. (c) The relative abundance of each isotopologue was extracted. (d) Calculated isotopologue populations at enrichments of ^2^H ranging from natural abundance to 6%. Combinations of these populations at varying relative proportions are compared to the experimentally measured data until the combination which minimises the residual between the experimental and calculated is found. The relative proportion of the non-natural abundance populations estimates the proportion of newly synthesised peptide over the labelling period.

The incorporation of ^2^H into protein was measured by observing shifts in the relative abundances of isotopologs of identified peptides. MS^2^ identification of MS^1^ features in unlabelled samples were transferred onto retention time aligned labelled samples allowing the extraction of the relative abundances of isotopologs of known elemental composition from plants grown in the day or the night (Figure 1b). This labelling strategy produced more subtle shifts in the isotopic envelopes than previous studies eg. 100% ^15^N labeling but were sufficient for reproducible measurement (Nelson et al. 2014; Li et al. 2017). MS scan information collected +/- 2 seconds from the elution peak of each peptide was averaged together to create a relative isotopolog profile (Figure 1c). The shape of this envelope for a peptide of known elemental composition is modified by the enrichment of ^2^H in the amino acid pool and the ratio of labelled to unlabelled peptide present. Interpretation of these shapes was tackled through iterating combinations and proportions of enriched and natural abundance populations to minimise the residual between the modelled and experimental data through the non-negative least squares regression algorithm as demonstrated previously (Nelson et al. 2014; Li et al. 2017) (Figure 1d).

Non synthetic incorporation of deuterium into proteins through hydrogen – deuterium exchange reactions has been shown to occur. These exchanges occur rapidly in sidechains, slowly into the amine backbone and not-at-all into non-exchangeable positions. Specifically optimised conditions are required to preserve these exchanges through bottom up proteomic sample preparation (Konermann et al. 2011). The low concentration of ^2^H in the labelling media used here, short labelling period and extensive opportunity for back exchange following removal of ^2^H from sample solvents during denaturation, proteolysis, fractionation, storage and analysis combined with not taking steps to preserve exchanged ^2^H effectively removed the possibility of deuterium exchange contributing to the labelling observed. Back-exchange of hydrogen for metabolically incorporated ^2^H likely does occur in exchangeable positions, however, the absolute numbers of ^2^H atoms present and the relative enrichment into each amino acid pool is not directly relevant to the new:old assessment made here; only that new protein has significant incorporation of heavy isotopes above natural abundance levels.

This analysis provided information on the ratio of peptide synthesised during the labelling period to that which was synthesised before it. To determine synthesis rates, the size of the protein pool at the beginning and end of the labelling period was also required. The size of each protein pool was tracked through a linked quantitation experiment in which unlabelled end of night and end of day samples were spiked 1:1 with a fully ^15^N labelled proteome reference mix. This mix was derived from plants grown in parallel in ^15^N labelled media from seed. This workflow allowed the determination of the ratio of newly synthesised to pre-existing protein from progressively labelled samples and determination of relative pool size changes from fully labelled spike in samples for each protein. These measurements were then weighted by the absolute abundance (Wang et al. 2015) and total protein amount per rosette to give a measurement of absolute synthesis and degradation rate per rosette in the light and dark during a 24 hour diurnal cycle.

Peptide abundance and isotope incorporation data was assembled into protein lists in two ways. The first generated incorporation data for each replicate and assembled a final protein measurement from the average of the three replicates. Due to stochastic variation in data dependent mass spectrometry acquisition, a smaller set of proteins could be reported on with this methodology than making a final protein measurement by assembling peptide measurements from across the three replicates directly. The direct assembly resulted in protein synthesis and degradation rates for approximately 350 proteins (Supplemental Table S1). While 350 is still numerically a small proportion of the total number of proteins encoded in the Arabidopsis genome, due to the abundance-based selection of data dependent mass spectrometry, these 350 proteins represent approximately 40% of the total protein amount in mature rosette leaves according to molar ratio estimates derived from paxdb (pax-db.org) (Wang et al. 2015). This methodology was performed in biological triplicate and each replicate consisted of the rosettes of at least four plants. Twelve hours of 5% ^2^H_2_O treatment had no noticeable effects on plant phenotype or leaf growth rate during the following days (Figure S1). This finding is consistent with previous studies in which low concentrations of ^2^H_2_O have been shown to have minimal impact on plant growth (Sideris, Mukherjee, and Vomvoyanni 1975, Yang et al. 2010; Zachleder et al. 2018).

Having derived synthesis rates for each protein relative to its pool size and tracked relative pool size for each protein over the diurnal cycle, Pax-db molar ratio data was used to define the size of each pool relative to total protein content (Wang et al. 2015). This series of calculations yielded a synthesis rate for each protein on a per total protein amount basis ie K_s_ per gram of protein. Total protein content for Arabidopsis rosettes at 21 days has previously been determined to be 2% of fresh weight (Li et al. 2017). As water content in young plants varies substantially over the day / night cycle an average fresh weight across all samples was used. Absolute synthesis and degradation rates were derived from these figures and a calculation of the total amount of ATP required to sustain this level of protein turnover made. These measurements assumed 5.25 molecules of ATP was required for each peptide bond in synthesis and 1.25 ATP molecules was required for the degradation of each bond (Kaleta et al. 2013; Pal et al. 2013). These figures indicate that for the proteins measured here, approximately three times more peptide bonds are formed in the day than are formed at night (Figure 2a). A day to night incorporation ratio has been measured previously at the total protein pool level through ^13^CO_2_ labelling (Ishihara et al. 2015). The ^13^C incorporation measurements were made through isolation of total protein before acid hydrolysis and isotope ratio determination of amino acids through GC-MS. Both methods produce a similar three to one ratio of day to night protein synthesis rate.

**Figure 2.**
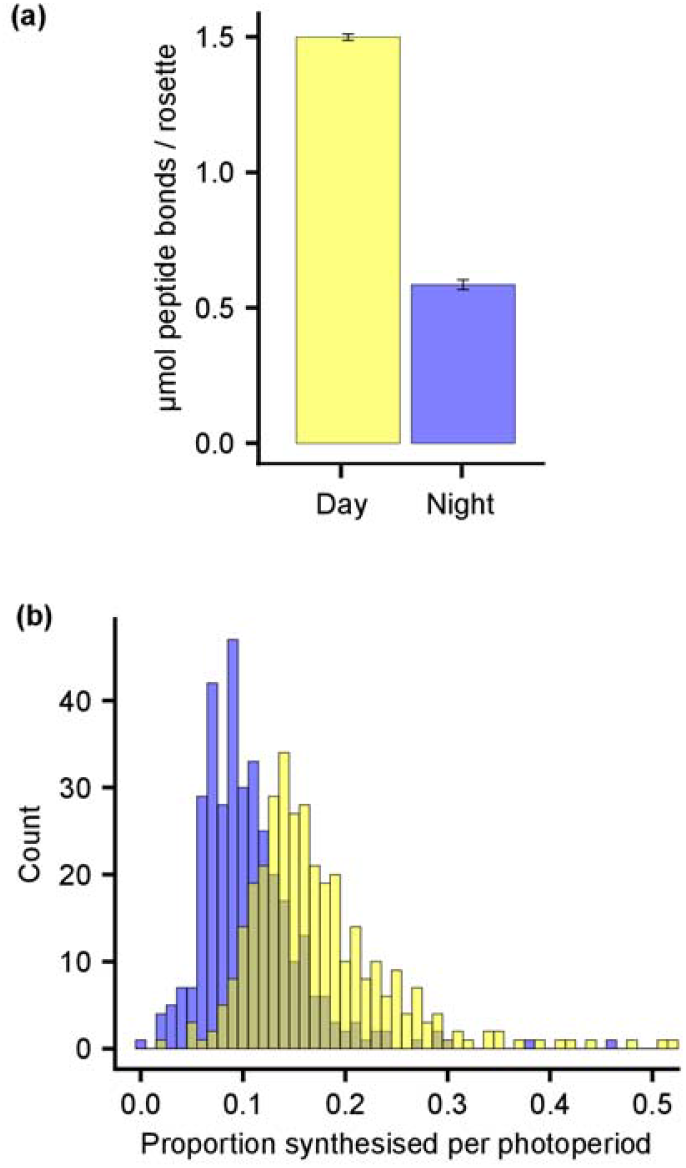
Pool level changes to protein synthesis rates in the day and night. (a) The total number of peptide bonds required to sustain the protein synthesis rates observed through incorporation of ^2^H were calculated on a per rosette basis, error bars are standard error. (b) Distribution of proportions of each protein pool synthesised during 12 hours of day (yellow) or 12 hours of night (blue).

The significant advantage of the ^2^H approach presented here is the ability to determine specific synthesis rate differences between a variety of proteins. We investigated the proportion of each protein pool synthesised in the day and the night and found that both were positively skewed distributions with standard deviations larger than the median values underlying a 53-fold range in night time and 42-fold range in day time values (Figure 2b, Supplemental Table S1). The mean proportion of each protein synthesised in the day was 19.3% while it was 11% at night (mean difference 8.26%, p 5.4e^-25^, t test). The higher skew in the day distribution indicated that day and night synthesis programs were not analogous even if shifts in the mean were discounted. We investigated this difference by comparing the positions of both functional categories and individual proteins within the distributions between the day and night.

### Day and Night Translational Program Trends

Separating the 350 proteins whose turnover rates were measured in both the light and the dark into functional categories showed that the magnitude of the reduction in synthesis rate at night was not uniform (Thimm et al. 2004). Although translation rates of all of the categories measured were significantly faster in the light (t test, p <.05), categories such as redox homeostasis, respiration and protein modification (1.4, 1.4 and 1.3 fold higher in light) showed smaller day/night differentials than photosynthesis, amino acid metabolism and protein synthesis (2.3, 3.0, 1.8 fold higher in light) (Figure 3a). The observed divergence in day to night synthesis rate ratios between functional categories suggests that regulation of translation rates is a specific, regulated process. Gross changes in substrate availability or general mechanisms of translation inhibition should suppress or promote translation rates uniformly, instead, these results suggest that translational regulation is more selectively controlled. The abundance of proteins at the functional category level either did not change or changed very little over the diurnal cycle. Of the categories with more than ten members in the 350 proteins measured, only redox homeostasis and protein synthesis show significant differences in their mean abundances (redox: fold change 1.09, pval 0.0089, protein synthesis: 1.039, pval 0.01) (Figure 3b). These results indicated that the changes in protein synthesis and degradation observed over the diurnal cycle were largely balanced, with both reducing at night.

**Figure 3.**
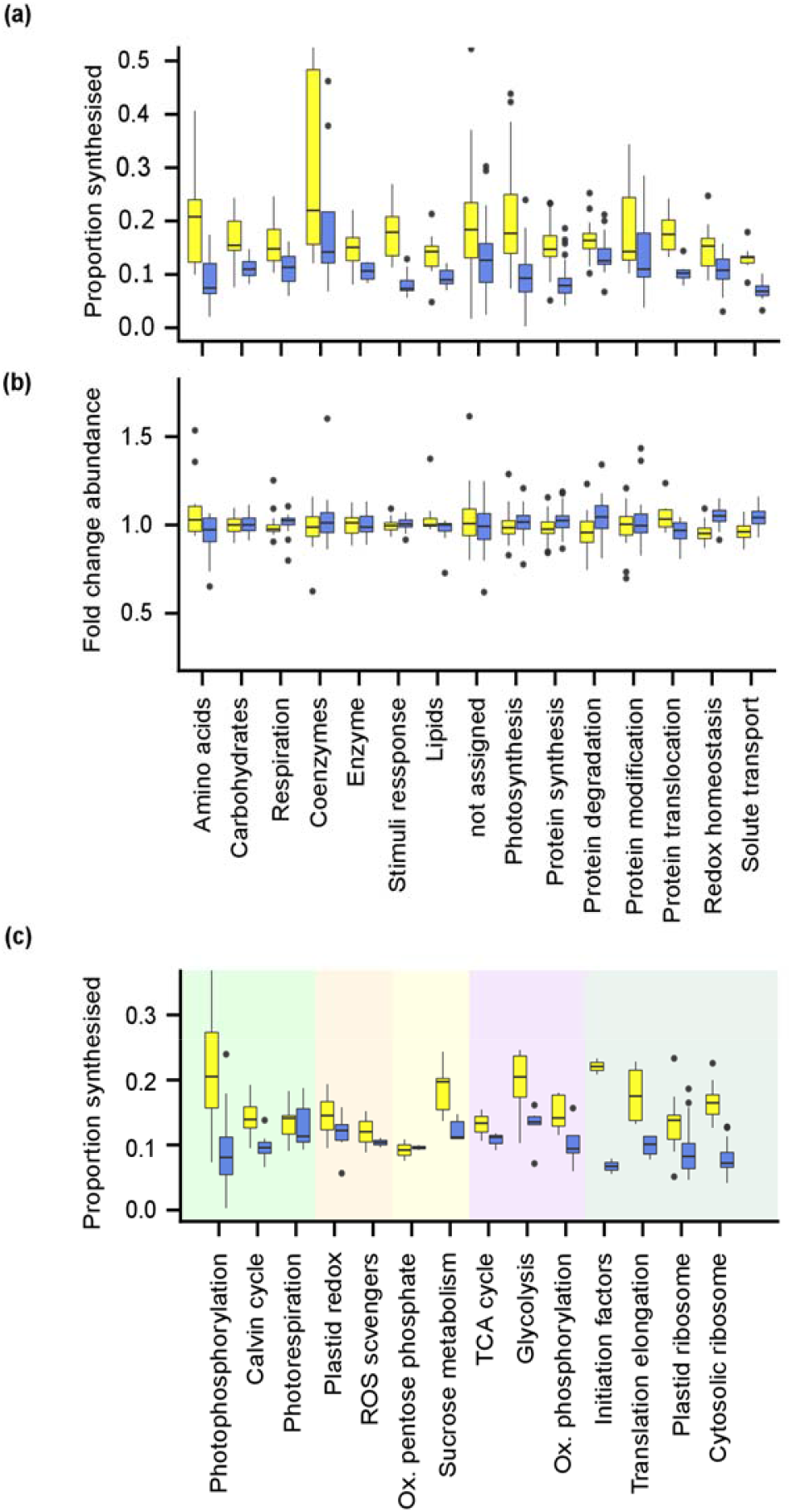
Functional categories of the newly synthesized Arabidopsis leaf proteins in the day and night. (a) Distribution of the proportion of each protein synthesised in 12 hours of day (yellow) or 12 hours of night (blue) grouped by the Mapman bin. Bins in which ten or more members of the category were observed are presented. (b) Distribution of the fold change in abundance of each protein from start of day to the end of the day (yellow) or start of night to the end of the night (blue] (c) Proportions of protein synthesised in selected subcategories of the Mapman bin system. Green background covers sub-bins which are members of the photosynthesis category. beige: redox homeostasis, yellow: carbohydrate metabolism, purple: cellular respiration, dark green: protein biosynthesis.

The Mapman bin system (Schwacke et al 2019) offers a hierarchical structure in which broad functional categories are broken down into specific pathways, complexes and subunits. We examined a selection of bins to gain a pathway level understanding of how translation is prioritised over the diurnal cycle (Figure 3c). Photophosphorylation showed more than double the translation rates during the day hours compared to night hours whereas supporting functions such as Calvin cycle enzymes, photorespiratory enzymes, plastid redox regulators and ROS scavenging proteins showed more similar translation rates in the day and night. Other categories which showed small differences in day / night ratios included well characterised pathways with important night-time functions such as oxidative phosphorylation, TCA cycle and ROS scavenging. Aligning with our finding that overall translation rates were lower at night, synthesis of initiation factors, elongation factors and cytosolic and plastid ribosome components were also found be comparatively lower at night.

### Day and Night Specific Translational Program Costs

Fold changes in day to night synthesis ratios ranged from 86 fold faster in the light - CHLOROPHYLL A/B BINDING PROTEIN 3 (CAB3) to 2.5 fold faster in the dark - PLASMA-MEMBRANE ASSOCIATED CATION-BINDING PROTEIN 1 (ATPCAP1). As a group, the light harvesting complex proteins showed the largest day/night synthesis rate differential with all members measured being at least ten times faster in the light. Of the proteins with higher translation at night, a number belonged to core functions of mitochondrial metabolism; SERINE HYDROXYMETHYLTRANSFERASE 1 (1.3 fold), CYSTEINE SYNTHASE C1 (1.3 fold), MANGANESE SUPEROXIDE DISMUTASE 1 (1.2 fold) and MALATE DEHYDROGENASE 1 (1.1 fold). However, both ends of the translation spectrum were dominated by plastid proteins. Those more rapidly translated during the day were largely photosynthetic machinery and those whose translation was night biased had more diverse functions; carbohydrate metabolism, lipid metabolism and REDOX maintenance.

In terms of the cost in ATP for the synthesis of each protein measured here, RBCL, subunits of Photosystem II and RUBISCO activase were the most expensive individual protein investments both in the day and night, though their relative ranking changed between conditions (Figure 4a,b). While the magnitude of the day/night synthesis rate variation was large, the overall ranking of ATP investment in different proteins was well conserved between day and night (Spearman correlation 0.91). Therefore, the notable large movement of specific proteins within these ranks is suggestive of diurnal regulation of translation rates either through transcript abundance and loading or subsequent regulation. Detailed lists of rank changes are shown in Supplemental Table S2. LHC proteins showed the largest change in cost ranks - CAB3 shifted 254 rank positions in the set of 350 proteins. A number of regulatory proteins also showed large shifts to a higher rank in the day; COLD, CIRCADIAN RHYTHM, AND RNA BINDING 2 (CCR2) - a known circadian cycling protein moved 101 ranks (Carpenter, Kreps, and Simon 1994), TRANSLATIONALLY CONTROLLED TUMOR PROTEIN 2 (TCTP2) - a nucleotide exchange factor in TOR signaling moved 96 ranks (Berkowitz et al. 2008), At1g13930 - an unknown protein previously identified in a circadian entrainment QTL (Darrah et al. 2006) shifted 78 ranks. Other proteins shifted higher in the night ranks; NADPH-DEPENDENT THIOREDOXIN REDUCTASE C (NTRC) - a central integrator and regulator of REDOX metabolism in chloroplasts (Spínola et al. 2008) and regulator of starch utilisation via modulation of AGPase (Michalska et al. 2009) shifted 49 ranks. Proteins with functions in lipid metabolism increased the most ranks at night: PATELLIN2, responsible for mediating the exchange of lipids between membranes shifted 133 ranks higher at night and ENOYL-ACP REDUCTASE 1, a member of the fatty acid synthase complex shifted 123 ranks higher.

**Figure 4.**
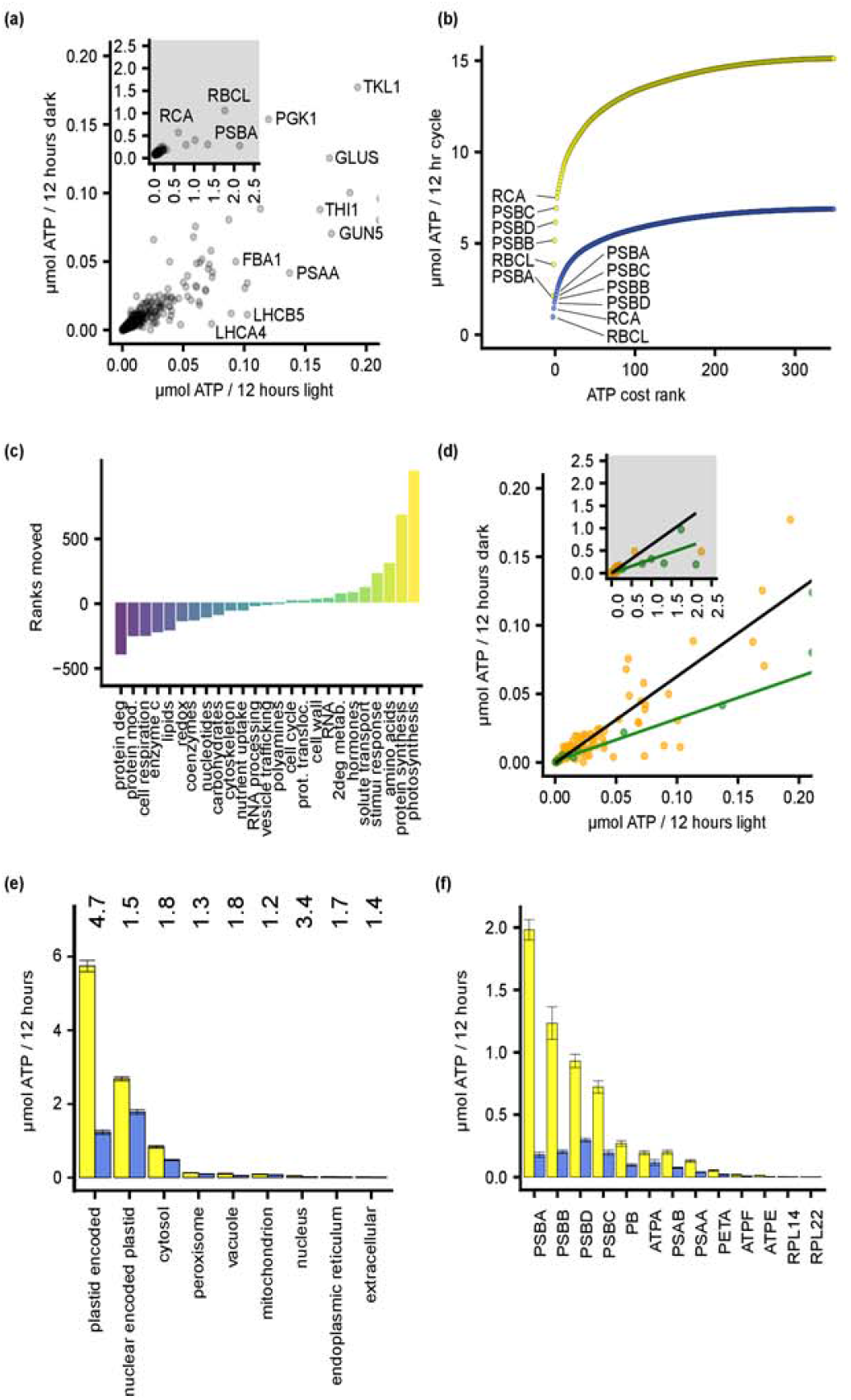
ATP cost of synthesizing different Arabidopsis proteins and functional categories of proteins in the day and night. (a) Scatter plot of the ATP cost to achieve the protein synthesis and degradation observed per rosette in the day and the night. The inset plot is an uncropped version highlighting the highest cost proteins in the day. (b) The cumulative total of ATP required for protein synthesis and degradation per protein in the day and the night. Proteins were arranged in descending order of ATP cost in the day (yellow) and night (blue). (c) Proteins were ordered in descending order of ATP cost in the day and in the night and the total number of rank positions moved per functional category were summed per Mapman category. Positive numbers refer to higher total rank in the day, negative to higher total rank at night. (d) Comparison of the ATP cost of plastid proteins encoded in the nucleus and translated in the cytosol (orange) or encoded and translated in the plastid (green) between the day and the night. Inset plot is an uncropped version of the main plot highlighting the five highest cost proteins in the day. Regression lines show linear models fitted to the cytosolic translated plastid dataset in black and the plastid translated set in green. (e) ATP cost of synthesis and degradation of measured proteins grouped by their subcellular localisations in the day and the night. Numbers at the top indicate the ratio of these averages. (f) ATP cost of each plastid encoded protein measured in the day and the night. Abbreviations: RCA rubisco activase, RBCL ribulose-1,5-bisphosphat carboxylase/oxygenase (RuBisCo) large subunit, PSBA photosystem II reaction center protein A, PSBB photosystem II reaction center protein B, PSBC photosystem II reaction center protein C, PSBD photosystem II reaction center protein D, PB ATP synthase subunit beta, ATPA ATP synthase subunit alpha, ATPF ATP synthase complex membrane CF0 subcomplex subunit b, ATPE ATP synthase complex peripheral CF1 subcomplex subunit epsilon, GLUS glutamate synthase, PSAA photosystem I subunit A, PSAB photosystem I subunit B, PETA photosynthetic electron transfer A, RPL14 plastidial ribosome large subunit proteome 14, RPL22 plastidial ribosome large subunit proteome 22, PGK1 phosphoglycerate kinase 1, TKL1 transketolase 1, GUN5 genomes uncoupled 5, FBA1 fructose-bisphosphate aldolase 1, LHCB5 light harvesting complex of photosystem II 5, LHCB4 light-harvesting chlorophyll-protein complex I subunit A4

Grouping the rank shift data by functional categories shows segregation of metabolism by diurnal cycle (Figure 4c). The photosynthesis category shifted the most in day translation rank, moving a total of 925 ranks between its members. Protein synthesis and amino acid metabolism categories were also preferentially translated during the day while protein degradation, modification and cellular respiration were preferentially translated at night. We investigated whether translation occurring in the plastid was differentially affected by the diurnal cycle when compared to cytosolic translation. To do this we compared the day:night ratio of translation rates of plastid proteins encoded in the nucleus vs those encoded in the plastid genome (Figure 4d). Translation on the plastid ribosome was found to be proportionally slower than cytosolic translation at night. To determine whether this effect was specific to the plastid ribosome or to the plastid itself we compared the ratio of day to night translation rates by organellar localisation and translation apparatus (Hooper et al. 2014) (Figure 4e). Most categories showed a day to night ratio of between one and two with only plastid translated and nuclear localised proteins falling outside these ratios. In terms of ATP cost per rosette the plastid translated proteins incur a far greater cost at ~6 *μ*mol ATP vs < .05 *μ*mol ATP for measured nuclear proteins. When day and night ATP costs of plastid encoded proteins are viewed individually, the reaction centre subunits of photosystem II stand out as having much greater reductions in ATP costs at night (Figure 4f).

In an effort to distinguish other proteins whose translation rate was controlled over and above the global shifts in translation rates observed, we normalised the median K_s_ of the three day and night replicates. By setting the median K_s_ to be equal between the day and the night, proteins with significantly different K_s_ may be considered to be subject to specific translational regulation over and above mechanisms which affect global translation rates. Student’s t tests showed 75 out of the 143 proteins which were present in all six replicates (three day, three night) had significantly different means at p < 0.05. After multiple test correction, (Benjamini and Hochberg 1995) 30 were below the corrected p < 0.05 to suggest evidence of specific translational regulation as a contributor to controlling the diurnal energy budget (Supplemental Table S3). This analysis again highlighted PSII reaction centre proteins and some nuclear-encoded light harvesting complex proteins which were found to have significantly slower protein synthesis rate at night. These proteins are exceptionally expensive proteins to synthesise and maintain having both high abundances and high degradation rates. Proteins translated faster at night than global averages included a number of stromal isoforms of glycolytic enzymes previously demonstrated to play central roles in the provision of substrates for respiration at night (Muñoz-Bertomeu et al. 2009).

### Correlation of Estimates of Protein Translation

Estimates of protein translation rates are commonly made by measuring the association of specific transcripts with one or more ribosomes or assumed to be well correlated with transcript abundance. As measurements of aggregate translation are still comparatively rare data, we compared protein translation rates inferred through polysomal loading studies and mRNA abundance with the translation rates measured here through stable isotope incorporation. We identified three studies focused on diurnal cycles which assessed either polysomal loading or transcript abundances on a day / night cycle. The first study quantitatively compared the translation state of specific transcripts by calculating a weighted sum of a transcript’s abundance in polysomal fractions vs the non polysomal fraction at two time points in the light and two in the dark in a 16h light / 8 h dark cycle (Missra et al. 2015). As that study uses a different day length to ours, we also included transcript abundance data from a 12 hour day / 12 hour night study by Bläsing et al. and a smaller set of polysomal loading data focused on enzymes of central carbon metabolism in a 12h light / 12h dark cycle by Piques et al. (Bläsing et al. 2005; Piques et al. 2009).

Missra et al. (2015) derived a protein synthesis rate by modelling diurnal variations in transcript abundance and translation state as sine waves before multiplying them together. As the processed data wasn’t available, we made a simple equivalent by calculating point to point area under the curves for day and night-time transcript abundance and translation state and multiplied them together to yield an estimate of aggregate translation. Comparing these ratios showed that the variation in polysomal loading of all transcripts cycled by much smaller magnitudes than aggregate translation would suggest (Figure S2a). Expanding the comparison to studies which were conducted with the same day length yielded better correlations but the magnitude of day to night cycling was consistently underestimated. The transcript abundance measurements made by Blasing et al. showed a slight positive relationship (Figure S2b) but only the 2009 study by Piques and colleagues which used QRT-PCR to quantify transcripts bound to polysomes for a select range of central carbon metabolic enzymes showed a strong positive relationship to ^2^H derived aggregate translation measurements (Figure S2c). While low correlations may in-part be explained by differences between labs and experimental setups, the metabolic changes between night and day are not subtle and clear positive correlation of the total datasets might reasonably be expected.

## Discussion

In this study we measured the amount of protein synthesis and degradation in the day and night for each of several hundred proteins. We tracked synthesis and degradation through the incorporation of ^2^H into newly synthesised protein while monitoring the degradation rate of the pre-existing unlabelled pool. These data differ to previous studies as they provide a direct measure of aggregate translational output rather than a proxy of ribosome association, so both global and transcript specific regulation is incorporated into the measurements. These data also employ an isotope which is abundant in peptides and is rapidly assimilated into amino acids independently of carbon fixation or nitrogen assimilation. This approach has allowed shorter time courses of labelling for protein specific measurements than has been previously possible. Measurements of aggregate translation distil the combined effects of regulation occurring at the genetic, transcript and translational levels and don’t rely on a range of assumptions made in calculating and reporting RNA based proxies for translation. This approach thus provides a different perspective to that generated through static measurements of abundance of transcripts and proteins.

### Leaf protein translation and degradation is highly diurnally regulated

Previous studies have shown that the rate of protein synthesis at the whole pool level in plant leaves is faster during the day hours. This effect paces starch usage, allowing plants to avoid carbon starvation at the end of the night (Sulpice et al. 2014) and delayed growth the following day (Gibon, Bläsing, et al. 2004). The high Spearman correlation of translation rate ranking between the day and the night, with overall lower rates at night agrees well with notion of general translational regulation promoting translation during the day and supressing it at night (OLeary et al. 2020; Merchante et al. 2017;Moore et al. 2016). This is also reflected by the bulk of the proteins measured here (~60%) moving less than 30 ranks in translation order between the day and the night (Supplemental Table S2).

### Specific examples of regulation of translation

In order to explain the remaining diversity of day:night translation rate ratios observed (Figure 2), gene or regulon specific rate modification mechanisms need to be evoked (Muench, Zhang, and Dahodwala 2012). Approximately 50% of the proteins assessed showed evidence of specific translational regulation as judged by their mean translation rate being significantly different after median day and night translation rates were equalised. We have attempted to assess the relative importance of general vs specific translational regulation in maintaining the diurnal energy budget by comparing the total cost in ATP over the 24 hour cycle consumed maintaining proteins which are apparently regulated by general mechanisms of translational control (89 of 143 proteins, 2.8 μmol ATP/rosette) vs those which shift translation rates significantly more than the median value (54 of 143, 8.9 μmol ATP/rosette). The large difference in ATP for synthesis cost of the specifically regulated set reflects the make-up of this pool; proportionally more high abundance, rapidly turning over proteins; the careful regulation of which would be likely to contribute to vegetative and reproductive fitness.

At the top of this list of ATP cost intensive, specifically diurnally regulated proteins are members of the PHOTOSYSTEM II reaction centre. The D1 subunit PSBA and its carefully regulated translation, degradation and assembly have been well studied (Anoman et al. 2016;Kato et al. 2015;Adir et al. 2003;Hutchison et al. 1996). Between PSBA, PSBB, PSBC and PSBD, approximately 4.9 μmol ATP per rosette are used in synthesis and degradation during the day, whereas 0.87 μmol ATP per rosette is used at night. Compared to total synthesis and degradation costs - 15.1 μmol per day and 6.9 μmol per night, the specific reduction of synthesis and degradation of these four proteins represents about half of the difference in protein translation and degradation costs making their specific control a significant contributor to the reduction in protein maintenance costs at night.

The selective translational regulation of light harvesting proteins has also been previously studied. The translation of these proteins occurs in the cytoplasm but is regulated in response to photosynthetic flux (Petracek et al. 1997) as well as light (Floris et al. 2013). As demonstrated in other species, individual members are also specifically translationally regulated; LHCBM is specifically regulated in response to shifts from low light to high light (McKim and Durnford 2006) when compared to LHCB4 in *C. reinhardtii*. This regulation has been explained mechanistically through the action of NAB1, a cytosolic RNA binding protein which selectively interacts with *LHCBM* transcripts and sequesters them away from the translational machinery (Mussgnug et al. 2005). Our data show that this phenomenon is evident in aggregate translation rate data but is not evident in mRNA association data (Figure S2a). The discovery that as many as 50% of the proteins whose translation rates were measured here show evidence of specific translational regulation suggests that such mechanisms may be more widespread than is currently appreciated.

The above examples illustrate proteins which are translated more slowly at night than mean rates would suggest. Plastid localised glycolytic enzymes however show the opposite trend: night time translation rates are significantly higher than the mean. The plastid isoforms of FRUCTOSE-1,6-BISPHOSPHATASE, GLYCERALDEHYDE 3-PHOSPHATE DEHYDROGENASE A-2 and PHOSPHOGLYCERATE KINASE all show higher translation rates at night than would be expected. The importance of plastid glycolysis in starch break down has proven difficult to assess due to the presence of transporters capable of exchanging glycolytic intermediates with the cytosol. These transporters have limited the utility of knock-out plants to assess the relative importance of cytosolic and plastid localised isoforms of glycolytic enzymes. The identification of phenotypes of knockouts of the plastid localised isoforms in specifically heterotrophic tissues (Anoman et al. 2016) suggests that these enzymes play important roles in non-photosynthetic energy provision in some tissues. The observation here that their night-time translation rates were specifically enhanced is supportive of a broader role in night time energy provision in photosynthetic tissues as well.

Selective changes in diurnal protein degradation rate are somewhat less studied. Circadian clock regulators are specifically degraded in timed daily patterns by the proteasome using a combination of phosphorylation and E3 ubiquitin ligase activation in plants (Fujiwara et al. 2008; Harmon, Imaizumi, and Gray 2008; Ooijen et al. 2011). There is scant evidence for selective diurnal degradation mechanisms in plants beyond core or peripheral clock components, albeit a wide variety of E3 ligases are being linked to their selective substrates in plants and many show developmental or stress responsiveness (Vierstra 2009; Shu and Yang 2017). Broader studies in mice liver have shown that daily oscillations in macro- and chaperone-mediated autophagy and proteasomal activity, effectively concentrates liver proteolysis during the day (Ryzhikov et al. 2019). To our knowledge, we lack evidence of how selective diurnal translation and degradation of the same components are coupled to generate the homeostasis observed in protein abundances (Figure 2b). It is only in much longer shifts to short- and long-day environments that the leaf standing proteome is modulated, and then only mildly so in most cases (Seaton et al. 2018).

### Divergence of transcript abundances and translation rates over the diurnal cycle

While there are a variety of global reports of diurnal variation in polysome association and Riboseq analysis (Bläsing et al. 2005; Missra et al. 2015), we noted a generally low correlation between these changes and aggregate translation rates. Most notably, an order of magnitude difference in the range of these transcript ribosome association measures and protein translation rate differences (Figure S2, Supplemental Table S1). We propose that reconciliation can come from recognition that different things are being measured. Where both polysome loading and translational output have been measured, significant discrepancies are seen to result. For example, Pal et.al. showed that while polysome loading in the night was approximately 2/3 of that during the day, the resulting translation was only 1/3 of the rate seen in day hours (Pal et al. 2013).

The polysome status of a transcript (how many ribosomes are associated with a specific transcript) is a general proxy for translation of any transcript based on the concept that as more ribosomes bind to a transcript, its rate of translation increases proportionally. Riboseq improves upon this technique by differentiating actively translating ribosomes from paused ribosomes (Calviello and Ohler 2017). Both of these techniques however are static snapshots of ribosome ‘input’. A number of assumptions lie between this input measurement and a direct measurement of translation. Particularly, that the rate of initiation of translation is rate limiting, and that the rate of polypeptide chain extension is constant across transcripts. Regulatory mechanisms such as codon usage, translation substrate inhibition, ribosome exit tunnel interactions and secondary mRNA structures may not be well accounted for when translational rates are inferred (Merchante, Stepanova, and Alonso 2017). In contrast, the quantitation of aggregate translation described in this report provides a collation of all inputs and processing constraints to provide the effective sum output of the ribosome.

### Protein translation correlates with pathway flux

While protein abundance is largely unchanged for the major metabolic and biosynthetic machinery of the leaf over the diurnal cycle (Figure 3b), the translational rate appears to correlate in part with the flux through metabolic pathways. For example, the strong day time translation of photosynthetic and amino acid synthesis machinery and indeed the protein synthesis machinery itself is linked to high rates of all three of these processes in the light. While some of this machinery is plastid-encoded and thus its rate of synthesis will be influenced by the lack of chloroplast ATP synthesis in the dark, a larger number are nuclear-encoded and reliant on the cytosolic ATP pool and machinery for translation. While this correlation could be put down solely to regulated use of ATP when it is more available in the light, it could also be that regulatory processes have hard-wired a biophysical link between use and the need for replacement. The specific case of the D1 subunit of PSII is an exemplar of this effect with damage and need for replacement being linked to catalytic mechanism and an 11-fold difference in turnover rate observed here (Supplemental Table S1). However, this effect could be broader, we recently highlighted the concept of catalytic cycles till replacement being a property of enzymes due to metabolic interactions and damage during operation (Tivendale et al. 2020). The high number of catalytic cycles in the light for some enzymes would therefore warrant a faster replacement schedule in the light to maintain enzyme pools at maximal enzyme capacity. When maximal extracted activities for 24 such enzymes were measured across diurnal cycles (Gibon, et al. 2004) it was found that many fell in maximal activity in the first few hours of the morning and rose across the day, only to fall again during the night. The often >1.5 fold diurnal variation in these reported enzyme activities (Gibon, et al. 2004) is not apparent in changes in protein abundance, suggesting that a combination of enzyme activation and deactivation is partially responsible and this is likely to be a combination of post-translational modifications of enzymes and damaged enzymes awaiting replacement.

## Conclusion

To match the recent focus on developing new high-throughput approaches to map and quantify different critical parameters affecting the inputs to translation, such as RNA structure, protein– RNA interactions and ribosome occupancy at the genome level (Merchante 2017), a renewed enthusiasm toward studying translation output is also warranted. Here we reinvigorate the potential for ^2^H labelling of plants to contribute to this process. In the detail we highlight a range of components for which input and output measures of translation differ markedly, pointing to selective translational regulation events that are currently hidden from view. In overview we show how the aggregate nature of these data allow insight into how limited energetic resources are prioritized to optimize metabolism and molecular fitness during day and night cycles in leaves.

## Materials and Methods

### Plant Growth Conditions

Arabidopsis thaliana Col-0 seeds were sown on 0.8% agar / ddH_2_O (w/v) contained in 2 ml Eppendorf tube lids punctured with ~3mm diameter holes. Lids were then placed into 24 tube racks and vernalised before transferring to 12 / 12 hour day (220 μmol/m^2^/second white light) night cycle at a constant 22°C. 0.5 strength Hoagland’s media was used throughout with solutions being changed weekly. Reagents including heavy isotopes were purchased from Sigma 5% ^2^H_2_O supplemented solutions were made up v/v with 99.9% deuterium oxide while ^15^N solutions contained K^15^NO_3_ 98% and ^15^NH_4_^15^NO_3_ 98% obtained from Sigma Aldrich.

### Sample preparation and analysis

Samples were prepared as detailed previously (Duncan et al. 2017). Resuspended peptides were fractionated into 96 samples by high pH reversed phase chromatography (Wang et al. 2011) before pooling into 12, each of which was analysed over 90 minutes (2-27% acetonitrile) on a Thermo Fisher Orbitrap Fusion with 75 μm x 500 mm c18 EASY-Spray column driven by a Dionex UltiMate 3000 RSLCnano system. Raw files were converted to *.mzML format (MSConvert, ProteoWizard 3.0.20079) and matched against Araport11 peptide sequences (www.arabidopsis.org) using CometMS (2019.01 rev. 5)(Eng, Jahan, and Hoopmann 2013). Results were rescored and protein lists assembled with PeptideProphet (Keller et al. 2002) and ProteinProphet (Nesvizhskii et al. 2003) (Trans-Proteomic Pipeline v5.0.0). MS^1^ feature lists were assembled with Dinosaur (1.1.0) (Teleman et al. 2016) and results inserted into PostgreSQL databases with MSFTBX (Avtonomov, Raskind, and Nesvizhskii 2016). Peptide identifications were matched to MS^1^ features in the R environment for statistical computing (R Core Team, n.d.) and retention times aligned by locally estimated scatter plot smoothing (LOESS). The primary mass spectrometry data files are available via ProteomeXchange with identifier PXD023274.

### Progressive isotope incorporation analysis

For approximately 120,000 peptide identifications per sample, a single isotope profile was assembled by extracting ten ion chromatographs - mono to plus 9 heavy from +/- 2 seconds around the maximum value of the monoisotopic peak and summing the intensities across this range using XCMS (Smith et al. 2006). In the case of ^2^H_2_O labelled samples, a correction factor for the retention time of labelled peaks due to the hydrophobicity modification caused by ^2^H incorporation was computed on a subset of high quality data but found to be negligible for the low incorporation seen here. Relative Isotope incorporation data termed labelled peptide fraction was calculated as detailed previously (Nelson et al. 2014) and updated (Li et al. 2017) with modifications detailed below. Data was filtered by requiring the the correlation between the isotopic envelope extracted from the unlabelled sample to the calculated natural abundance envelope to be greater than 0.97, The intensity apex of the associated MS^1^ feature be greater than 50,000, that the deviance of the NNLS result be less than 0.009, mono isotopic mass errors were zero and the heaviest NNLS population was less than 3%. A protein group list was assembled by analysing all unlabelled runs as a single sample which allowed consistent assignment of peptides to protein groups across the experiment. Data which passed the above criteria were ordered by the sum of intensities of the MS^1^ feature and a mean labelled peptide fraction was derived from all available measurements of the top 3 peptides of different sequence for each protein group. This assembly was performed both on a per replicate basis and a total data set basis in which the data from all replicates were pooled. Each protein group had a minimum of 3 different peptide sequences and a minimum of 5 independent measurements of labelled peptide fraction to be included in the total data set or 3 different peptide sequences with at least 3 independent measurements to be included in the distinct replicate analysis. Each protein group had an average of 8.84 independent measurements of 3.9 different peptide sequences in the total data set and 3 different peptide sequences with 3 independent measurements in the replicate analysis. The standard deviation and relative standard deviation for each protein group in both assemblies was calculated and groups with RSD < 0.25 or SD < 0.06 were retained for analysis.

### Relative quantitation analysis

Samples were taken from plants grown in parallel in both unlabelled and ^15^N media. In the case of the ^15^N labelled plants samples were taken at the end of night and end of day before being mixed together 1:1 on a fresh weight basis prior to protein extraction. Unlabelled plants were sampled at end of night and end of day and protein extraction preformed separately. Unlabelled samples were mixed 1:1 with the labelled protein mixture (amido black) before digestion. Determination of the labelled to unlabelled ratio was performed as above with the following modifications - correlation of natural abundance envelope to theoretical was not used to filter the data and the average ratio of natural abundance to heavy was normalised to 0.5 on a per replicate basis.

### Leaf area analysis

Pots containing 24 plants were grown on a 3 x 3 grid as above were photographed hourly by triggering a Cannon EOS600D with an Aputure AP-TR1C. Each pot of 24 plants was aligned on a 3 x 3 grid and images post processed by dividing into 9 using Imagemagick’s crop and repage functions (The ImageMagick Development Team 2020) and total green pixels (rgb 210,210,20) per image counted using a 15% ‘fuzz’ factor.

## Material distribution footnote

The author responsible for distribution of materials integral to the findings presented in this article in accordance with the policy described in the Instructions for Authors (www.plantcell.org) is: A. Harvey Millar

## Supplemental Data

Figure S1. Relative leaf area measurements of plants grown with and without 12 hours of 5% ^2^H_2_O.

Figure S2. Comparison of aggregate protein synthesis rate regulation with diurnal transcript and polysomal occupancy levels.

Table S1. Summary table of protein synthesis and degradation rates in Arabidopsis leaves during the night and day

Table S2. Summary table of changes in rank of night to day protein synthesis rates in Arabidopsis leaves

Table S3. Summary of protein synthesis rate data from night and day in Arabidopsis leaves following median normalisation of replicates

## Acknowledgments

This work was supported by the Australian Research Council awards CE140100008 and DP180104136 (to A.H.M.). Analysis for this work was performed by the WA Proteomics Facility as a node of Proteomics Australia and was supported by infrastructure funding from the Western Australian State Government in partnership with Bioplatforms Australia under the Commonwealth Government National Collaborative Research Infrastructure Strategy.

## Author contributions

OD and AHM designed the research. OD performed the experiments and data analysis, OD and AHM wrote the paper.

